# Learning Protein Representations with Conformational Dynamics

**DOI:** 10.1101/2025.10.06.680789

**Authors:** Dan Kalifa, Eric Horvitz, Kira Radinsky

## Abstract

Proteins change shape as they work, and these changing states control whether binding sites are exposed, signals are relayed, and catalysis proceeds. Most protein language models pair a sequence with a single structural snapshot, which can miss state-dependent features central to interaction, localization, and enzyme activity. Studies also indicate that many proteins assume multiple, functionally relevant shapes, motivating approaches that learn from this variability. Here we present DynamicsPLM, a protein language model conditioned on ensembles of computationally generated conformations to derive state-aware representations. DynamicsPLM improves predictive performance across protein–protein interaction, subcellular localization, enzyme classification, and metal-ion binding. On a widely used protein–protein interaction benchmark, it achieves a four-point accuracy gain over the strongest baseline. On a curated test set enriched for proteins with multiple conformational states, the margin increases to eleven points. These findings argue for a shift from static to dynamics-aware modeling, in which conformational variability is treated as informative. By elevating conformational state to a central element of machine learning in protein biology, this work advances modeling toward mechanisms that better reflect how proteins operate in cells and provides a route to actionable hypotheses about when and how binding, signaling, and catalysis occur.^*^

## 1 Introduction

Many proteins exist not as a single rigid fold but as a shifting ensemble of conformations, transitioning among distinct functional states as they bind substrates, undergo post-translational modifications, or interact with partner molecules [5]. These changes in structure—whether subtle domain flexing or large-scale allosteric transitions—are critical to the protein’s role in signaling, regulation, and catalysis. In recent work, generative models, trained on protein structures, molecular dynamics results, and experimental data, have been employed to make inferences about molecular dynamics, yielding protein equilibrium ensembles [11, 13, 19].

Despite advances in predicting different configurations and equilibria of conformations in protein interactions, protein language models (PLMs) that underpin much of today’s structure-based prediction rely on static structural snapshots. As an example, a great deal of research assumes a single AlphaFold-predicted coordinate set [7] or Protein Data Bank (PDB) entry [1], overlooking that conformational heterogeneity can influence biological behavior [10, 12, 20]. This choice can introduce concrete inconsistencies in downstream features that are state-dependent, such as pocket accessibility, catalytic geometry, and interface exposure.

A motivating example is triosephosphate isomerase (UniProt P04789), whose active-site *ω*-loop (169–176) toggles between *open* and *closed* states. Open apo structures (e.g., PDB: 5TIM) admit substrate, whereas ligand-bound complexes adopt a closed loop that seals the pocket (e.g., PDB: 1NEY). The AlphaFold model [7] for this protein predicts a closed loop with high confidence (pLDDT *>* 85), despite the known open/closed ensemble of the protein. Conditioning models on this single prediction risks, conflating ligand-induced closure with intrinsic properties of the protein and can bias predictions on multi-conformation targets.

To address this limitation, we introduce *DynamicsPLM*, a PLM that accounts for structural dynamics by integrating over multiple conformations via a dynamic embedding layer. The model is fully compatible with generative conformation models (e.g., BioEmu [11]); their predicted conformations can be used directly as inputs to the dynamics embedding layer, without other changes in the pipeline. We treat the conformation generator as independent and frozen: its conformations serve as inputs to the dynamics embedding layer, while we train only the embedding layer and the PLM encoders. This design avoids perturbing the conformation generator and prevents conflating gains from improved generation with gains from our fusion mechanism.

As shown in Figure 1, the methodology departs from prior PLMs in its core representation. Rather than conditioning on a single structure, DynamicsPLM constructs an ensemble of plausible protein conformations. Each conformation is discretized into structure tokens from a 3D alphabet produced by a VQ–VAE–pretrained autoencoder [22]. These structure tokens are combined with amino acid tokens to form structure-informed sequences, which are then passed through our proposed dynamics embedding layer to yield per-residue, weighted, conformation-aware token embeddings.

**Fig. 1:**
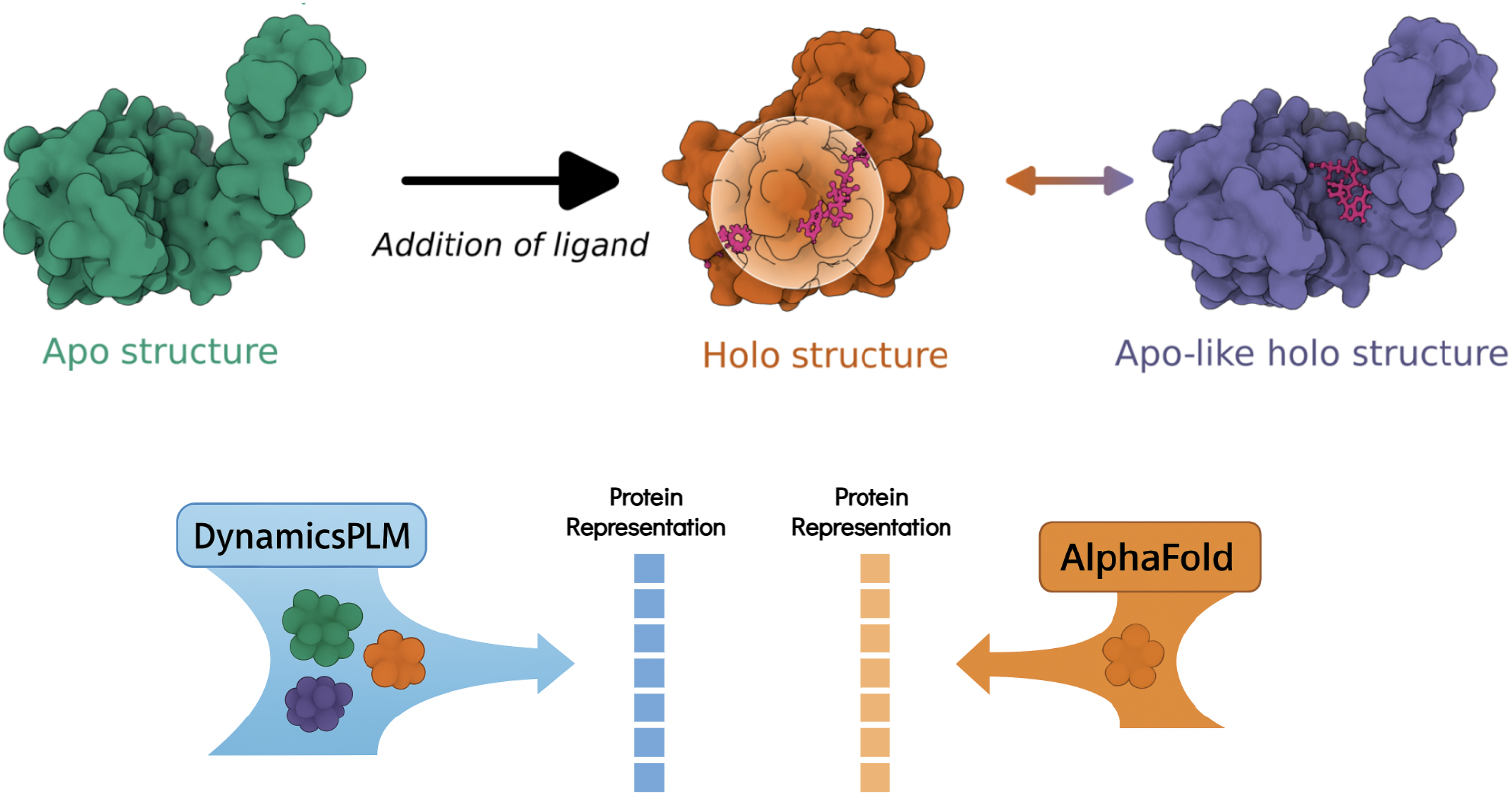
Conformational change enables interaction. *Top:* Conformation-dependent binding pocket of adenylate kinase [3]: an *apo* conformation (left; PDB 6S36, green), ligand-bound *holo* conformation with a formed/expanded pocket (center; PDB 8CRG, orange), and an *apo-like holo* conformation that retains an apo-like global fold while accommodating the ligand (right; PDB 6F7U, purple). *Bottom:* Illustration of DynamicsPLM protein embedding that is built via conditioning on ensembles of conformations and learning state weights that emphasize binding-competent configurations, while preserving alternatives. In contrast, single-structure pipelines (e.g., SaProt [20]) rely on AlphaFold-derived models [7] and use a single, static structure—often a holo-like conformation (*≈* 70% of cases) [17], thus, missing ensemble effects that are critical for interaction-dependent function.

We find that the resulting representation is *sensitive to both sequence context and conformational flexibility*, allowing the model to learn structure-conditioned embeddings that reflect the probabilistic nature of proteins. This design improves the model’s ability to resolve paralogs—proteins with near-identical sequences and the same overall fold but differing in the amplitude or occupancy of local motions (e.g., loop closures, hinge bends), and to identify functionally relevant regions defined not by static geometry but by their propensity for dynamical shifts.

A straightforward alternative is to encode several conformers separately and then average their embeddings. However, *such averaging collapses distinct structural modes into a single vector that corresponds to no single realized physical conformation*, muting signals tied to specific functional states (e.g., open versus closed loops, or ion-bound versus unbound pockets). In contrast, DynamicsPLM represents distributions of states rather than averages of states: for each residue, we construct a distribution over discrete structural microstates inferred from a conformational ensemble. This distribution approximates the local conformational free-energy landscape and preserves multi-modality, allowing the encoder to prioritize the state most relevant to the task at hand. We present a comparison (Supplementary Section S4.2) that contrasts our method with simple averaging of embeddings across generated conformers, which yields lower performance while incurring additional computational overhead.

We evaluate DynamicsPLM on several fundamental protein tasks: HumanPPI (human protein-protein interactions), metal ion binding, enzyme commission (EC), and subcellular localization prediction (DeepLoc). We ensure strict separation of close homologs by verifying that every protein shares at most ≤30% pairwise sequence identity with proteins in other splits. Across test sets, DynamicsPLM consistently outperforms the strongest baseline, improving accuracy by +4.0 points in HumanPPI, +1.9 in metal ion binding, and +1.6 in DeepLoc, and increasing EC *F*_max_ by +0.9 points. All gains are statistically significant (*p <* 0.05, two-tailed paired *t*-test with Holm–Bonferroni correction). These gains are significant because the evaluated tasks underpin many real-world applications, including drug discovery, which relies on accurate modeling of protein-protein interactions, and the elucidation of disease mechanisms through subcellular localization prediction.

To assess sensitivity to conformational dynamics, we re-evaluate all methods on proteins with multiple experimentally determined conformations from the CoDNaS-Q database [4]. In this protein dynamics subset, DynamicsPLM shows larger improvements over the strongest baselines: +11.1 points in HumanPPI, +3.85 in metal ion binding, +6.25 in DeepLoc, and +6.5 points in EC *F*_max_. All gains are statistically significant (*p <* 0.05, two-tailed paired *t*-test). These results reinforce our central claim: Conditioning on conformational ensembles, rather than a single static snapshot, yields the greatest benefits for proteins confirmed experimentally to have pronounced structural heterogeneity. In contrast, improvements are modest when conformational dynamics are limited. For proteins with only a single experimentally determined structure, performance remains comparable to strong baselines, suggesting that ensemble conditioning does not introduce noise in the absence of conformational diversity.

Taken together, these results indicate that conditioning on conformational ensembles, rather than a single snapshot, yields more reliable, biologically grounded protein representations and should become a default strategy for structure-informed PLMs in applications ranging from interaction and localization prediction to drug discovery.

## 2 Results

### 2.1 Overview of DynamicsPLM

DynamicsPLM is a broadly applicable PLM (see Figure 2) that accounts for structural dynamics by integrating over multiple conformations via a unique dynamic embedding layer. The model is trained on amino acid sequences paired with 3D structures predicted by AlphaFold2 [7]. In a first stage, a generative conformation model proposes multiple plausible conformations per protein. We utilize RocketSHP [19], a scale-efficient variant of BioEmu [11] shown to reduce prediction error, though the framework remains compatible with alternative generators. For each conformation, we build a structure-aware sequence by pairing residue tokens with discretized structural tokens produced by a VQ–VAE autoencoder [21]. The dynamic embedding layer then performs per-residue, cross-conformation fusion using ensemble-derived weights (normalized to sum to one), yielding a single sequence of conformation-aware token embeddings. The fused sequence is processed by transformer encoder layers initialized with pre-trained SaProt-650M weights [20] to produce a refined protein representation. Then, per downstream task, we incorporate task-specific classification heads to enable predictions for these tasks. See Section 4 for the full algorithmic details and implementation specifics.

**Fig. 2:**
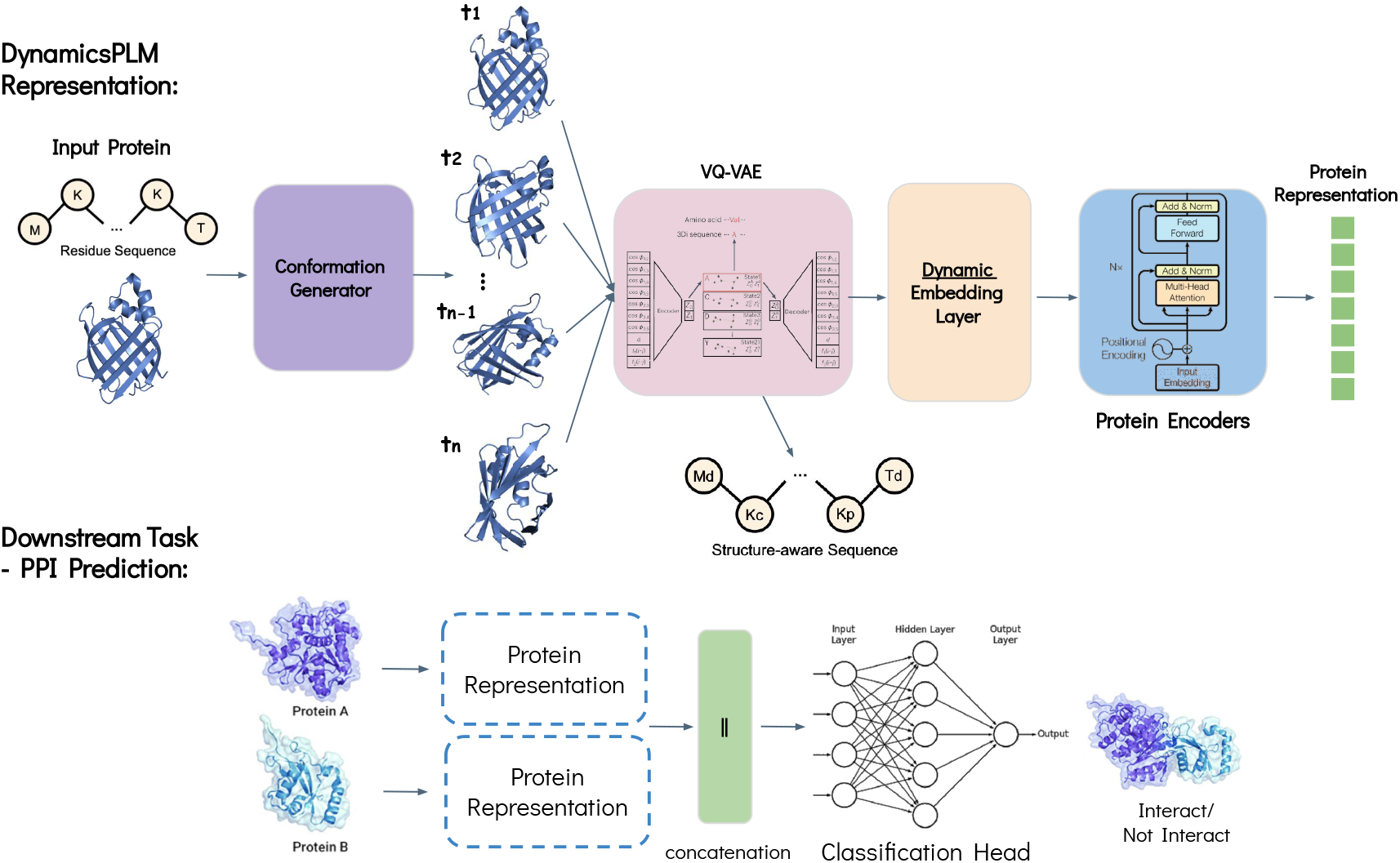
DynamicsPLM architecture. Top: The model is trained on amino-acid sequences together with 3D structures predicted by AlphaFold2 [7]. We obtain multiple plausible conformations per protein via a conformation generator. For each conformation, we construct a structure-aware sequence by pairing residue tokens with discretized structure tokens produced by a VQ–VAE–pretrained autoencoder [22]. The dynamics embedding layer then performs weighted, per-residue fusion across conformations estimated from token frequencies within the ensemble (normalized to sum to one), yielding a single sequence of conformation-aware token embeddings. The fused sequence is passed through the encoder layers to produce a refined protein representation. Bottom: Example of the downstream-task phase for the HumanPPI task, where the model predicts whether two given proteins interact.

### 2.2 Benchmark Tasks

We assess the effectiveness of our approach for protein representation learning on a diverse set of tasks, selected according to the top-performing joint structure–sequence PLMs, structure-only PLMs, and sequence-only PLMs [20, 23, 24]. These tasks span several biological fields, including protein-protein interaction prediction, localization prediction, and binding site classification (see Supplementary Section S1.2 for detailed task descriptions, and Supplementary Section S2.1 for evaluation and dataset details). To ensure the validity of the results, we validate all downstream task splits by enforcing that sequences in one set share no more than 30% Needleman–Wunsch sequence identity with any sequence in the other sets. For the HumanPPI task, we use group-wise splits by protein, assigning all interactions of a given protein to a single split to prevent partner leakage; the 30% identity filter is applied at the protein-cluster level prior to pairing. This homology filtering ensures minimal overlap between training and evaluation data, minimizing the risk of data leakage.

Benchmark test sets are strictly held out. To minimize pre-training exposure from the conformation generator, we screened all test proteins against the RocketSHP [19] pretraining corpus and removed any sequence with *≥* 30% identity to that corpus.

#### 2.2.1 Benchmark Results

In Table 1, we compare the performance of DynamicsPLM with six SOTA baseline methods (see Supplementary Section S1.1 for details) across diverse biological downstream tasks. DynamicsPLM significantly outperforms all baselines, achieving the best scores on every task. We compute Cohen’s *d* effect sizes [2] to quantify the magnitude of these improvements and observe large effects (*d >* 0.8) across the board. We attribute these gains to the inclusion of the dynamic modality.

**Table 1:**
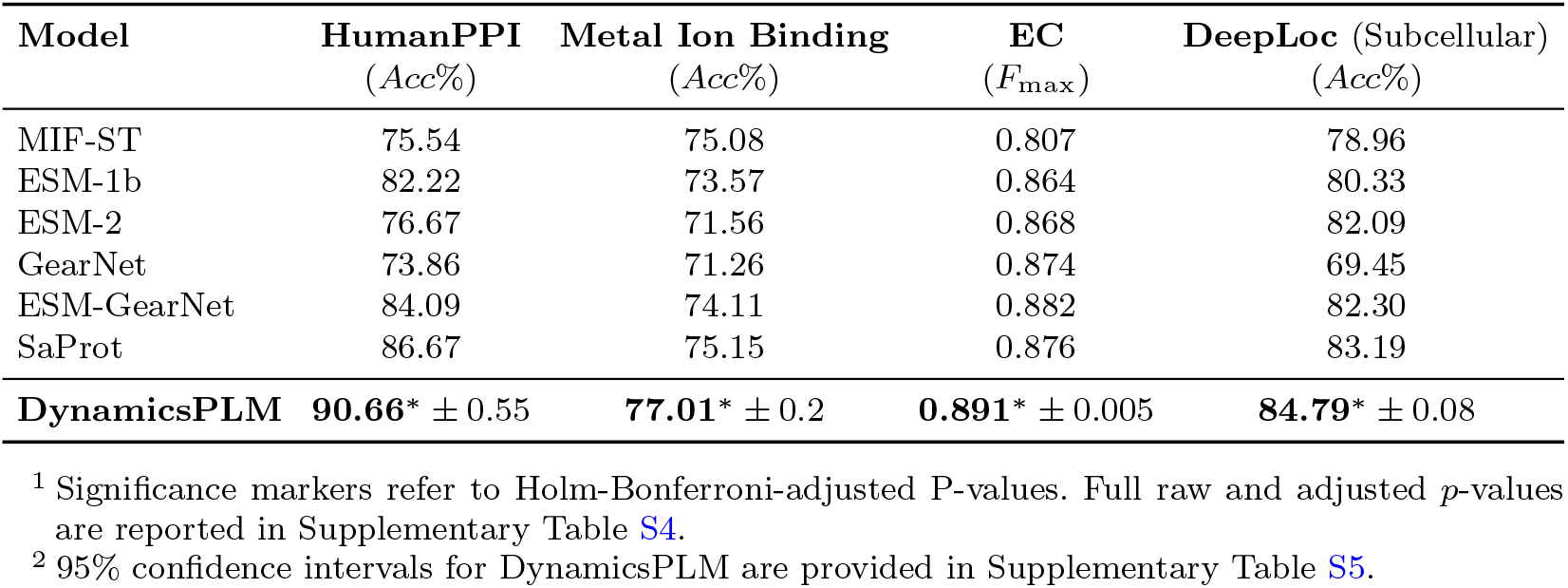
Experimental results on four downstream tasks. Statistically significant results (*p <* 0.05) using a two-tailed paired *t*-test across proteins in the test set (with Holm–Bonferroni correction across baseline comparisons for each task)^1^ are marked with an asterisk (*). For DynamicsPLM, we report the mean and standard deviation over 5 independent fine-tuning runs (different seeds; HumanPPI: *n* = 180, Metal Ion Binding: *n* = 665, EC: *n* = 1604, DeepLoc: *n* = 2747). The best result is highlighted in bold.

DynamicsPLM delivers the largest gains where state dependence is central and maintains an edge on tasks with weaker or implicit dynamic signals, highlighting ensemble-aware representations as a robust and generalizable foundation for protein function prediction. HumanPPI shows the largest gains, presumably because protein–protein interactions are strongly state-dependent [16]. Interfaces form or break as binding patches become exposed or buried, loops open or close, and allosteric shifts change local geometry. By conditioning on an ensemble of conformations rather than a single structure, DynamicsPLM captures these state-dependent features and their sequence determinants, yielding a four-point improvement compared to the strongest baseline [20] in this task. In practice, this leads to better detection of partners mediated by flexible loops and termini, as well as by transient, ligand- or metal-stabilized interface motifs, effects that existing models often overlook.

We find that Metal Ion Binding benefits from ensemble conditioning at known coordination sites, that DeepLoc gains reflect subtle state-linked exposure patterns relevant to trafficking signals, and EC improves despite limited explicit state annotations—suggesting that the learned representations generalize even when dynamic cues are only indirectly present in downstream labels. We further examine the performance gaps and find that ensemble context increases prediction confidence without sacrificing calibration. Confidence stratification and reliability diagrams across tasks (Supplementary Section S3.3) show upward shifts in accuracy at matched confidence and improved calibration slopes. Overall, these results underscore the value of incorporating conformational ensembles into protein modeling.

We conducted ablation studies across all benchmarks isolating key components of our model: (i) the weighting function (Supplementary Section S4.1), (ii) replacing the dynamic embedding layer with a mean-pooling operator (Supplementary Section S4.2), and (iii) swapping the conformation generator for an alternative (Supplementary Section S4.3).

#### 2.2.2 “Dynamic” Proteins

To quantify the impact of conformational heterogeneity on downstream performance, we repeated the evaluation on a filtered subset of each task’s original test set (see Table 2). The subset includes only “dynamic” proteins, identified using CoDNaS-Q [4], a curated dataset of proteins with multiple experimentally determined structures (i.e., conformers) that quantifies backbone variability across the ensemble. For each protein, CoDNaS-Q provides pairwise backbone deviations (using *Cα* RMSD differences), enabling us to select cases with non-trivial conformational diversity. Exact filtering criteria and per-task subsets are detailed in Supplementary Section S2.2.

**Table 2:**
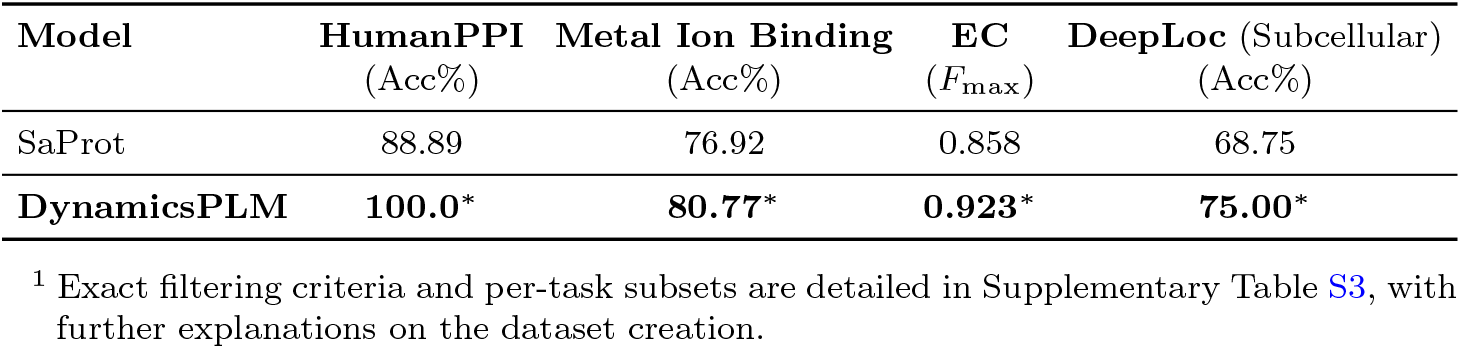
Experimental results for SaProt and DynamicsPLM on four downstream tasks, evaluated on a filtered subset of each task’s original test set. The subset includes only “dynamic” proteins^1^, identified by their known conformational ensembles and pairwise *Cα* RMSD differences based on the CoDNaS-Q dataset [4]. Statistically significant results (*p <* 0.05) using a two-tailed paired *t*-test across proteins in the test set are marked with an asterisk (*). Sample sizes after filtering: HumanPPI (*n* = 18), Metal Ion Binding (*n* = 101), EC (*n* = 134), DeepLoc (*n* = 16). The best result is highlighted in bold.

| Model | HumanPPI<br>(Acc%) | Metal Ion Binding<br>(Acc%) | EC<br>($F_{\max}$) | DeepLoc (Subcellular)<br>(Acc%) |
| --- | --- | --- | --- | --- |
| SaProt | 88.89 | 76.92 | 0.858 | 68.75 |
| **DynamicsPLM** | **100.0\*** | **80.77\*** | **0.923\*** | **75.00\*** |

<sup>1</sup> Exact filtering criteria and per-task subsets are detailed in Supplementary Table S3, with further explanations on the dataset creation.

DynamicsPLM outperforms SaProt [20], the strongest baseline, on every benchmark with statistically significant gains, consistent with the expectation that state-dependent biology benefits from ensemble conditioning. The results in the dynamic subset mirror the trends in the entire test set, but the gains are markedly larger. Taken together, these findings underscore the strength of DynamicsPLM on proteins that sample multiple conformations, widening its margin over the existing PLMs.

The largest improvement appears on HumanPPI, consistent with the main benchmark, with a gain of +11.11 points (100.0% vs. 88.89%). Other tasks benefit as well, with improvements of +3.85 points on Metal Ion Binding, +6.5 on EC, and +6.25 points on DeepLoc; all differences are significant (*p <* 0.05).

The dynamic-subset analysis isolates the value of ensemble conditioning: when proteins are known to sample multiple states, predictions improve markedly across tasks. Practically, these findings support three recommendations. First, when multi-conformer data (experimental or molecular dynamics derived) are available, ensemble-aware encoders should be preferred. Second, benchmarks and evaluation protocols should preserve conformational diversity rather than collapsing to a single representative structure. Third, simple triage rules (e.g., RMSD thresholds or bound/unbound flags) can identify cases where ensembles are most likely to yield benefits.

### 2.3 Curated Experimental Case Studies

To underscore the design and practical relevance of DynamicsPLM, we examine model outputs on a curated subset of human protein pairs with prior *in vivo* or *in vitro* experimental evidence for protein–protein interactions [15]. We present two case studies that highlight complementary error modes addressed by ensemble conditioning: recovering a state-dependent true interaction and (ii) suppressing a biologically implausible pair. In each case, we compare DynamicsPLM with the top-performing single-structure PLM baseline [20] to show how conformational ensembles reveal binding-competent interfaces that a single conformer may miss. We also evaluated a recently reported (2025) interaction [18] to test whether DynamicsPLM captures state-dependent interfaces and can predict interactions beyond our historical datasets.

#### Interacting pair: ATG10—ATG7

ATG10 (UniProt: Q9H0Y0) is an E2-like conjugating enzyme and ATG7 (UniProt: O95352) an E1-like activating enzyme in the ATG12*→*ATG5 autophagy pathway. Structural studies indicate that ATG7–ATG10 recognition is state dependent, with noncanonical E1–E2 contacts and loop repositioning between unbound and bound forms [8, 9]. The ground truth label for these proteins is a confirmed interaction. DynamicsPLM predicts interaction with probability (0.7903), whereas the single-conformer baseline (i.e., SaProt [20]) predicts non-interaction (wrong label) with a high probability of (0.600). We infer that evaluating multiple plausible conformations allows the model to capture binding-competent arrangements—surfaces on ATG7 and ATG10 that become complementary only in certain states, which may be under-represented by any single structure.

#### Non-interacting pair: MDM4—GCSAM

MDM4 (UniProt: O15151) is a nuclear regulator of p53 with a peptide-binding N-terminal pocket and a C-terminal RING domain that heterodimerizes with MDM2; multiple partner-specific states have been described [14]. GCSAM, also known as HGAL (UniProt: Q8N6F7), is a germinal-center B-cell adaptor localizing to the plasma membrane and cytoplasm, with ITAM/SH2-binding motifs that recruit kinases and modulate B-cell receptor signaling [6]. Here, the ground truth is non-interaction. DynamicsPLM predicts non-interaction with probability (0.7469), while the single-conformer baseline (i.e., SaProt [20]) predicts interaction (wrong label) with high probability (0.8972). We infer that scanning across conformers did not reveal a solvent-exposed, conformer-consistent interface on MDM4 compatible with a membrane adaptor such as GCSAM, and the ensemble model accordingly lowers the score.

#### Interacting pair: TMEM9—CLCN3 (recent, membrane-state dependent)

A recent study (2025) reports that TMEM9 (UniProt: Q9P0T7) directly binds and inhibits the endosomal chloride-proton exchanger CLCN3 (UniProt: P51790) [18]. Cryo-EM resolved the TMEM9–CLCN3 complex and showed that PI(3,5)P_2_ stabilizes the interface, indicating a membrane lipid–dependent, conformation-specific association. Cell-based assays in human cells corroborate the complex and its regulatory effect. On this pair, DynamicsPLM assigns a high interaction probability (0.7205), whereas a single-structure baseline predicts non-interaction (0.5963). We interpret this discrepancy as DynamicsPLM capturing multiple conformations—including lipid-stabilized, binding-competent states—that a single static structure under-represents, which is crucial for understanding this interaction.

#### Conclusion

DynamicsPLM provides informative variation: it elevates positives that appear only in specific states and suppresses negatives that lack a viable interface in any state. By conditioning on conformational ensembles, DynamicsPLM elevates state-dependent true interactions and suppresses pairs lacking a viable interface across states, improving performance. Because our cases are grounded in curated *in vivo* or *in vitro* evidence, the model’s high-confidence outputs translate into concrete, testable hypotheses. In practice, this helps prioritize experiments and reduce cross-compartment false positives, linking representation learning to bench-ready validation plans.

## 3 Discussion

We introduced DynamicsPLM, a structure-informed PLM that integrates information across conformational ensembles. Treating state as part of the representation improves prediction across interaction, localization, and enzyme-function tasks, with the largest gains where biology is demonstrably state dependent (e.g., protein–protein interfaces). As a core contribution, we introduce residue-wise ensemble conditioning that preserves multi-modal conformational occupancy via token histograms and learns to select function-relevant modes, outperforming encode-then-average baselines. In our evaluations, these gains are accompanied by more reliable confidence estimates, indicating improvements beyond simple threshold effects.

A practical attribute of DynamicsPLM is modularity. The protein conformation source can operate independently and be revised or replaced without architectural changes to the overall representation, enabling rapid adoption of improved generators.

The analysis of boosts in the conformational dynamics-subset clarifies when consideration of ensembles matters most: the gains in predictive power are seen precisely on proteins where there is independent evidence for multiple conformational states— where biology depends on proteins assuming distinct changes in structure. This suggests concrete guidance: It is valuable to harness ensemble-aware encoders when bound–unbound pairs or simple geometric criteria indicate conformational diversity, and avoid collapsing benchmarks to a single representative structure.

We note several limitations in the methodology and experiments. First, ground-truth state labels are scarce: for many proteins, it is unknown which conformations are biologically active under the conditions implied by the downstream labels. Consequently, the approach depends on an external conformer generator whose coverage and biases may vary across families. Second, the backbone RMSD used for screening is an imprecise surrogate for functional state—global motions can inflate RMSD without changing activity, while local rearrangements that gate binding may occur with little change in RMSD. Future work should expand dynamic benchmarks with curated experimental ensembles and explicit state annotations.

## 4 Methods

We introduce DynamicsPLM, a broadly applicable PLM that accounts for structural dynamics by integrating over multiple conformations via a unique *dynamic embedding layer*. The full implementation details, including computational complexity analysis, are presented in Supplementary Section S1.3.

### Structure-histogram vector

Let a protein *p* = (*S, R*) be defined by its amino acid sequence *S* = (*s*_1_, *s*_2_, …, *s*_*L*_) and its 3D structure *R*, where *s*_*i*_ *∈ V* denotes the *i*-th residue and *V* is the standard residue alphabet. Given the sequence and structure, a conformation generator produces *K* possible conformers 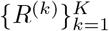.

Each conformer *R* is discretized into structure tokens of a 3D alphabet ℱ of size *M*, produced by a VQ–VAE–pretrained autoencoder [22], yielding per-residue structure tokens 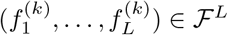.

Then, we compute for every position *i*, a structure-histogram vector:

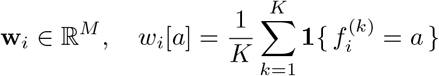

Where *w*_*i*_[*a*]*≥* 0*∀a ∈* ℱ and ∑_*a∈ℱ*_ *w*_*i*_[*a*] = 1.

This vector represents the empirical distribution over structure tokens at *i* across the ensemble. We refer to **W** = (**w**_1_, …, **w**_*L*_) as the SHP tensor (structure-histogram probabilities).

### Dynamic embedding layer

We maintain a learnable 3D embedding table **E** *∈* ℝ ^| *V*|*×M ×D*^ that contains a vector for every (amino acid, structure token) pair. We leverage the pre-trained SaProt-650M [20] embeddings to initialize this table, leveraging its strong contextual understanding of structure-aware sequences.

For residue *s*_*i*_ *∈ V* we form a structure-weighted embedding:

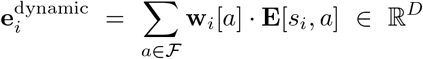

Let 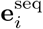 be the pre-trained embedding associated with the input token at *i* (i.e., an amino acid token). We fuse the two embeddings with a fixed convex combination to produce the final embedding, combining the sequential embedding with the dynamics one:

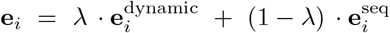

This aggregation yields per-residue, weighted, conformation-aware protein embedding **e**_*p*_ = (*e*_1_, …, *e*_*L*_) *∈* ℝ^*L×D*^.

### End-to-end pipeline

During training, each protein is passed through the dynamic embedding layer, producing a dynamic protein embedding *e*_*p*_. Then, multiple encoder layers produce a contextual protein representation *z*_*p*_ *∈* ℝ^*L×D*^, where *L* is the sequence length and *D* is the feature dimension. Then, after the encoder layers, task-specific classification heads are incorporated to enable predictions for the specific downstream tasks.

We initialize the encoders with a large pre-trained PLM, SaProt-650M [20], leveraging its strong contextual understanding. Using SaProt allows us to fairly isolate and assess the contribution of dynamics conformation integration. Alternatively, other advanced PLMs can also be used within the proposed framework.

The model is fully compatible with other generative conformation models (e.g., BioEmu [11]); their predicted conformers can be used directly as inputs to the dynamics embedding layer without any changes to the pipeline. We present such a replacement in the Supplementary Section S4.3.

We treat the conformation generator as frozen: its conformations serve as inputs to the dynamics embedding layer, while we train only the dynamic embedding layer and the PLM encoders. This design avoids perturbing the conformation generator and prevents conflating gains from improved generation with gains from our fusion mechanism.

To strengthen our key design choices, we present ablation tests examining the weighting function (Supplementary Section S4.1), replacing the dynamic embedding layer with a mean-pooling operator (Supplementary Section S4.2), and replacing the generative conformation model with an additional generator (Supplementary Section S4.3).

## Data availability

All input data used in this study are publicly available from the sources cited in the paper. Processed benchmark datasets for all tasks, including dynamic annotations and the exact train/validation/test splits, are available at https://github.com/kalifadan/DynamicsPLM.

## Code availability

The DynamicsPLM source code, pre-trained weights, and training/inference scripts are released under an open-source licence at https://github.com/kalifadan/DynamicsPLM. We also provide deterministic seeds, exact package versions, and instructions to regenerate the structural ensembles, ensuring full reproducibility of our results. Models were implemented in PyTorch (https://pytorch.org).

## Author contributions

D.K. led the research, developed the architecture and its training. E.H. and K.R. contributed technical advice and ideas. D.K., E.H. and K.R. wrote the paper.

## Additional information

### Supplementary information

The online version contains supplementary material available at Nature website.

## S1 DynamicsPLM Algorithm

In this section, we provide the full algorithm details of DynamicsPLM, a broadly applicable PLM that accounts for structural dynamics by integrating over multiple conformations via a unique dynamic embedding layer.

### S1.1 Baselines

Incorporating SOTA models as baselines and following SaProt [15] for fair comparison, we include ESM-1b [13] and ESM-2 [11], which are considered top-performing sequence-based models. Structure-based baselines include GearNet [21] and MIF-ST [19], while ESM-GearNet [20] serves as a representative joint sequence–structure model. We also evaluate against SaProt itself [15], the current leading PLM that incorporates AlphaFold-derived structure tokens.

### S1.2 Tasks

We provide a comprehensive overview of the downstream tasks, selected according to the top-performing joint structure–sequence PLMs, structure-only PLMs, and sequence-only PLMs [15, 20, 21], to rigorously evaluate the effectiveness of our proposed approach for protein representation learning.

#### S1.2.1 Protein-Protein Interaction Prediction

Reliable detection of protein–protein interactions (PPIs) is essential for deciphering cellular processes and uncovering therapeutic targets, especially when the interactions have not been previously characterized [8]. We use the HumanPPI dataset from the PEER benchmark [18] to evaluate binary interaction prediction between protein pairs. Following established best practices in this task [15, 18], we report accuracy as the primary evaluation metric.

#### S1.2.2 Protein Function Prediction

We evaluated protein function using the Metal Ion Binding [7] task, a binary classification task designed to predict the presence of metal ion binding sites within a protein, evaluated by accuracy, which is the common practice in this task [2, 15] for application usages.

#### S1.2.3 Protein Localization Prediction

We adopt the DeepLoc dataset [1], a subcellular localization task, including a 10-class multiclass classification task. We use accuracy as the primary performance metric following similar studies in this field [15].

#### S1.2.4 Protein Annotation Prediction

We evaluate protein functional annotation using the Enzyme Commission (EC) number prediction task from the DeepFRI benchmark [5]. This task is formulated as a multi-label classification problem, where each protein may be assigned one or more EC labels. *F*_max_ score is used for evaluation.

### S1.3 Implementation Details

We utilized the pre-trained SaProt-650M [15] as the base protein encoder, and its pre-training embedding to initialize our learned embedding table. The hyperparameters of DynamicsPLM (summarized in Supplementary Table S1) were selected via grid search based on performance on the validation set of the training dataset. As a result, the weight of the dynamic component in the embedding layer is set to *λ* = 0.5. Also, the number of conformations generated per protein is set to *K* = 20. The dimensionalities of the protein representations are set to *D* = 1280. To accommodate long protein sequences, inputs are truncated to a maximum of 1,024 tokens. All training is performed using mixed-precision arithmetic to improve memory efficiency and omputational throughput.

In addition, we incorporate task-specific classification heads to enable predictions of downstream tasks, following SaProt [15] settings, to ensure fair comparisons.

**Table S1:** Training hyperparameters for DynamicsPLM.

| Parameter | Value |
| --- | --- |
| Optimizer | AdamW |
| $\beta_1, \beta_2$ | 0.9, 0.98 |
| Weight decay | 0.01 |
| Learning rate | $2 \times 10^{-5}$ |
| LR schedule | Fixed |
| Warmup steps | – |
| Decay start / end | – |
| Training epochs | Task-specific |
| Batch size | 64 |

#### S1.3.1 Computational Complexity

All experiments were conducted on 4*×*NVIDIA A100 (80 GB) GPUs, with all run-time variables specified in our publicly available code. DynamicsPLM fine-tuning required approximately 24 hours (*∼*96 GPU-hours) per downstream task. The primary over-head scales with *K*, the number of conformations generated per protein, and with the depth of the encoder stack. In contrast, SaProt [15] was pre-trained from scratch for roughly three months on 64 GPUs. By reusing SaProt’s pretrained embeddings as initialization and integrating dynamic context via a lightweight dynamic embedding layer, DynamicsPLM achieves an approximate 99% reduction in training cost relative to full PLM pre-training.

At inference time, DynamicsPLM introduces only modest overhead. SaProt encodes at *≈* 0.014 seconds per 1,000 residues, whereas DynamicsPLM requires *≈* 0.016 seconds for the same input—an increase of about 12.5%. This overhead is minor, given the observed accuracy gains and full compatibility with standard PLM infrastructure. To avoid redundant computation, we generate conformations once per protein and cache them for reuse at inference.

**Table S2:** Benchmark dataset statistics for the benchmark tasks.

| Dataset | Category | Metric | Train | Valid | Test |
| --- | --- | --- | --- | --- | --- |
| HumanPPI [18] | PPI Prediction | Accuracy | 26319 | 234 | 180 |
| Metal Ion Binding [7] | Protein Function Prediction | Accuracy | 5067 | 662 | 665 |
| EC [5] | Protein Annotation Prediction | $F_{\max}$ | 13089 | 1465 | 1604 |
| DeepLoc (Subcellular) [1] | Protein Localization Prediction | Accuracy | 8747 | 2191 | 2747 |

## S2 Datasets Overview

### S2.1 Benchmark Datasets

Supplementary Table S2 summarizes the benchmark datasets, including task categories, evaluation metrics, and the sizes of training, validation, and test splits.

### S2.2 Dynamic Proteins Datasets

To isolate test cases where conformational state is most likely to affect function, we intersect each benchmark’s test proteins with CoDNaS-Q [4]—a curated resource that aggregates multiple experimentally determined conformers per protein and reports conformational diversity via the maximum pairwise *C*_*α*_-RMSD across conformers. We then retain proteins satisfying two criteria: (i) a minimum number of conformers and a non-trivial range of pairwise *C*_*α*_-RMSD (thresholds below). Requiring *≥*3 conformers (or *≥* 2 for DeepLoc, where coverage is sparser) stabilizes diversity estimates and follows prior practice of quantifying state variability across multiple structures rather than single pairs.

Our thresholds are informed by structural biology conventions and prior analyses [3, 12], combined with distributional statistics on our test proteins to exclude outliers. Very small backbone deviations are typically attributable to thermal/experimental fluctuation and are not interpreted as distinct states. Accordingly, we set a lower bound of 0.4 Å on the *C*_*α*_-RMSD range to exclude trivial noise while admitting subtle but meaningful changes.

Results are robust to small perturbations: varying RMSD thresholds by *±*0.1 Å and the conformer-count criterion by *±*1 changes subset sizes but leaves all qualitative conclusions unchanged (same direction of effects and significance). Importantly, we preserve the original dataset labels and splits; we only select test proteins that meet the criteria. Summary statistics are shown in Table S3.

**Table S3:**
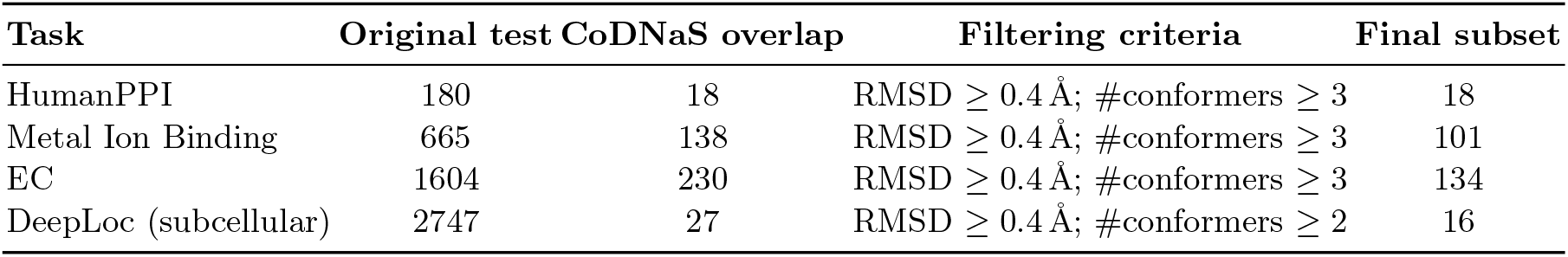
Dynamic-subset filtering per task. “CoDNaS overlap” counts test-set proteins present in CoDNaS-Q [4] before filtering. RMSD refers to the range of pairwise *C*_*α*_-RMSD across conformers.

## S3 Statistical and Reliability Evaluation

### S3.1 Statistical Significance Tests

We perform two-tailed paired *t*-tests across proteins in the test set to assess whether DynamicsPLM significantly outperforms each baseline for a given task. For each task, raw *p*-values (DynamicsPLM vs. each of six baselines) are adjusted using the Holm–Bonferroni procedure over that task’s baseline comparisons. In Supplementary Table S4 we list the raw *p*-values and report the maximum Holm–Bonferroni–adjusted *p* across those comparisons. Statistically significant differences (*p <* 0.05) indicate that the observed performance improvements are unlikely to have occurred due to stochastic variation.

**Table S4:**
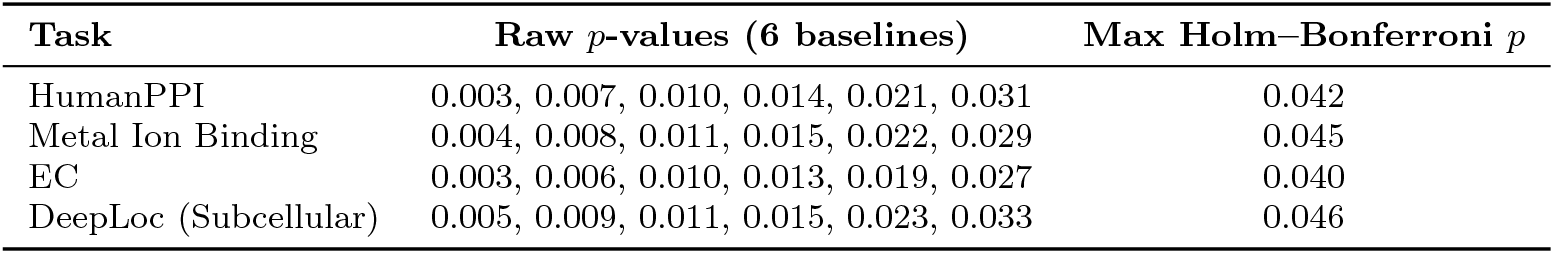
Raw and Holm–Bonferroni–adjusted *p*-values for DynamicsPLM across tasks. Raw *p*-values are from two-tailed paired *t*-tests across proteins. We report the maximum Holm–Bonferroni–adjusted *p* across these six comparisons for the task.

### S3.2 Confidence Intervals

To provide a clearer estimate of uncertainty across the five independent fine-tuning runs, we report 95% confidence intervals (CIs) for DynamicsPLM alongside the mean and standard deviation (Supplementary Table S5). Confidence intervals are computed using a two-tailed *t*-distribution with *n* = 5 (independent fine-tuning runs with different seeds).

**Table S5:** Mean ± standard deviation and 95% confidence intervals for DynamicsPLM across the benchmark tasks (five fine-tuning runs; different seeds).

| Task | Mean ± SD | 95% CI |
| --- | --- | --- |
| HumanPPI (Acc%) | 90.66 ± 0.550 | [89.98, 91.34] |
| Metal Ion Binding (Acc%) | 77.01 ± 0.200 | [76.76, 77.26] |
| EC ($F_{\max}$) | 0.891 ± 0.005 | [0.885, 0.897] |
| DeepLoc Subcellular (Acc%) | 84.79 ± 0.080 | [84.69, 84.89] |

### S3.3 Reliability Diagrams

As classification networks must not only be accurate but also reliable in their uncertainty estimates, calibrated confidence [6] is essential for interpretability and downstream decision-making. We evaluate and plot the expected calibration error (ECE) [6] for the classification tasks, using 10 equal-width bins over the confidence or predicted value range, following the calibration equation [6, 9]. DynamicsPLM consistently exhibits improved calibration over SaProt [15], with lower ECE in DeepLoc (0.086 vs. 0.143) and Metal Ion Binding (0.051 vs. 0.116). We observed similar trends in the additional tasks. Reliability diagrams (see Supplementary Figure S1) show that DynamicsPLM tracks the identity line more closely than SaProt across the confidence bins.

**Fig. S1:**
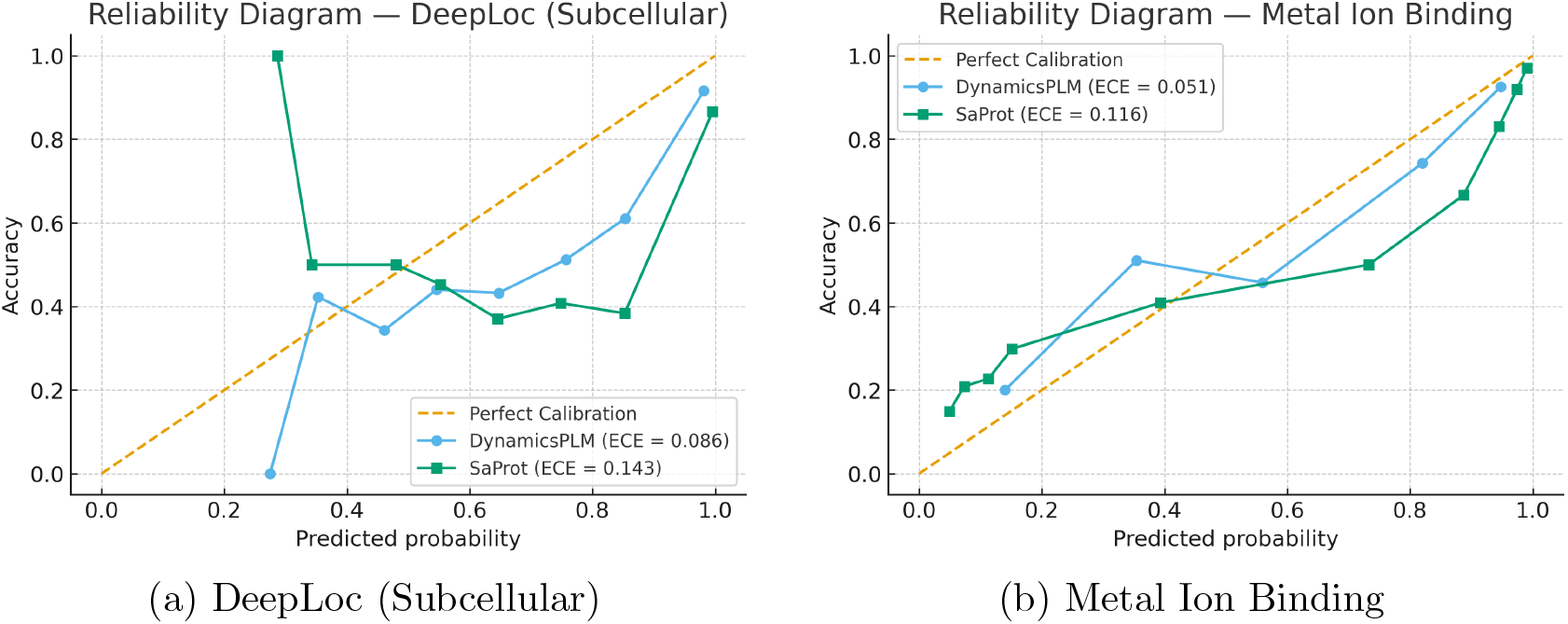
Reliability diagrams for DeepLoc (Subcellular) and Metal Ion Binding. DynamicsPLM (blue) demonstrates better calibration than SaProt (green) in both tasks. In these classification tasks, DynamicsPLM achieves lower ECE while closely following the perfect line.

## S4 Ablation Tests

### S4.1 Ablation: Fixed vs. Learned Mixing Weight *λ*

In the full DynamicsPLM, we keep the mixing weight *λ* fixed (shared across residues and inputs), which we found to be stable and to incur negligible overhead. As defined in Section 4:

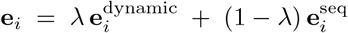

We ablate this choice by learning a per-residue *λ*_*i*_ *∈* [0, 1] via a lightweight gate (all other settings unchanged). The gate is implemented as `LayerNorm`(*H*) *→* `Linear`(*H, H*) *→* `GELU` *→* `Dropout`(0.1) *→* `Linear`(*H*, 1) *→* `Sigmoid`, where *H* is the embedding dimension; the final bias is initialized to *−*0.85 (so *σ*(*−* 0.85) *≈* 0.30) to conservatively favor sequence features. In all full-model experiments we set *λ* = 0.5.

Supplementary Table S6 shows that learning *λ*_*i*_ does not yield measurable gains and slightly increases run-to-run variance. Empirically, the gate converges toward a near-constant weight (low entropy across positions), indicating that a global trade-off between **e**^dynamic^ and **e**^seq^ suffices for these benchmarks; a fixed *λ* is therefore preferred for simplicity and stability.

**Table S6:** Ablation of the static–dynamic mixing weight. “Fixed *λ*” is the full DynamicsPLM; “Learned *λ*_*i*_” uses a per-residue gate. Statistically significant results (*p <* 0.05) using a two-tailed paired *t*-test across test proteins are marked with an asterisk (*). The best result is in bold.

| Task | Fixed $\lambda$ (full) | Learned $\lambda_i$ |
| --- | --- | --- |
| HumanPPI (Acc%) | **90.66\*** | 90.11 |
| Metal Ion Binding (Acc%) | **77.01\*** | 76.94 |
| EC ($F_{\max}$) | **0.891\*** | 0.887 |
| DeepLoc Subcellular (Acc%) | **84.79\*** | 84.24 |

### S4.2 Ablation: Mean-Pooling Over Generated Conformers

In this ablation, we replace the DynamicSHP layer with mean-pooling over the generated conformers. For each protein, we generate *K* conformers, similar to the dynamic embedding layer (see Section 4), and convert each to a structure-aware sequence. Then, we run the encoder once per conformer, and average the resulting embeddings across the *K* conformers using a mean-pooling operation.

Averaging full-sequence embeddings collapses multi-modal residue states into a single intermediate representation. Functionally distinct conformations (e.g., open vs. closed loop; ion-bound vs. unbound pocket) are blended, which can obscure the signal the classifier needs at decision time. In contrast, DynamicsPLM does not average conformers: it builds a per-residue distribution over structural tokens and feeds that distribution to the model. This retains multi-modality, closer to a local free-energy landscape, so the network can learn to emphasize the functional state when it matters instead of being forced into a single mean embedding. Even as *K →∞*, mean-pooling converges to an unstructured expectation over complete sentences that ignores residuewise uncertainty the model could exploit; our distributional embedding preserves that uncertainty explicitly at each position.

Empirically (see Supplementary Table S7), mean-pooling underperforms the full DynamicsPLM on all benchmarks and shows higher run-to-run variance. This supports exposing the positional distribution directly to the encoder rather than compressing it via pre- or post-encoder averaging.

Moreover, when compared against the top-performing single-structure baseline (SaProt), mean-pooling is equal or slightly worse, indicating that indiscriminate aggregation dilutes informative states and amplifies noise from outlier/non-physiological conformers. In contrast, DynamicsPLM’s selective integration of ensemble information preserves signal while maintaining lower variance, yielding consistent gains over the single-conformation baseline.

Finally, our method is computationally more efficient: it runs the encoder only once per protein, whereas mean-pooling requires *K* encoder passes. This yields an approximate *K* times reduction in wall-clock compute for the same *K*.

**Table S7:**
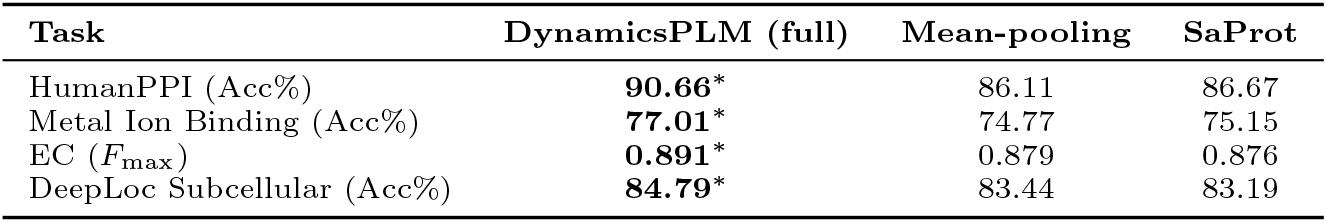
Ablation comparing our proposed DynamicsPLM to (i) a mean-pooling over *K* embeddings produced from generated conformers and (ii) a static-structure baseline (SaProt [15]; single conformation). Statistically significant results (*p <* 0.05) via a two-tailed paired *t*-test across test proteins are marked with an asterisk (*). The best result is in bold.

### S4.3 Ablation: Conformation Generator

To test whether gains arise from the fusion mechanism rather than generator-specific idiosyncrasies, we compare DynamicsPLM under two conformation generators— RocketSHP [14] and BioEmu [10]—while holding the encoder, data splits, training protocol, and ensemble size fixed (*K* = 10).

Across benchmarks (Table S8), BioEmu closely tracks RocketSHP yet is modestly lower on all tasks, without statistically significant differences; the preserved ranking and effect sizes indicate that ensemble-conditioned fusion, not properties of a particular sampler, drives the improvement. To mitigate concerns about leakage, we verified for BioEmu that no evaluation protein overlaps its pre-training corpus above 30% sequence identity.

These results mirror independent evaluations of structure–histogram probabilities (SHP) on ATLAS [17], where RocketSHP attains lower *K*-divergence [16] than BioEmu (1.089 vs. 1.923 at matched sampling budgets; lower is better) [14].

**Table S8:** Ablation comparing our proposed DynamicsPLM under two conformation generators (RocketSHP [14], BioEmu [10]) and a static-structure baseline (SaProt [15]; single conformation). Statistically significant results (*p <* 0.05) via a two-tailed paired *t*-test across test proteins are marked with an asterisk (*). The best result is in bold.

| Task | DynamicsPLM (RocketSHP) | DynamicsPLM (BioEmu) | SaProt |
| --- | --- | --- | --- |
| HumanPPI (Acc%) | **90.66\*** | **89.44\*** | 86.67 |
| Metal Ion Binding (Acc%) | **77.01\*** | **76.54\*** | 75.15 |
| EC ($F_{\max}$) | **0.891\*** | **0.883\*** | 0.876 |
| DeepLoc Subcellular (Acc%) | **84.79\*** | **84.46\*** | 83.19 |

## Footnotes

* Code, model weights, and inference scripts are available at https://github.com/kalifadan/DynamicsPLM.

